# Complete protection by a single dose skin patch delivered SARS-CoV-2 spike vaccine

**DOI:** 10.1101/2021.05.30.446357

**Authors:** Christopher L.D. McMillan, Jovin J.Y. Choo, Adi Idris, Aroon Supramaniam, Naphak Modhiran, Alberto A. Amarilla, Ariel Isaacs, Stacey T.M. Cheung, Benjamin Liang, Helle Bielefeldt-Ohmann, Armira Azuar, Dhruba Acharya, Gabrielle Kelly, Germain J.P. Fernando, Michael J. Landsberg, Alexander A. Khromykh, Daniel Watterson, Paul R. Young, Nigel A.J. McMillan, David A. Muller

## Abstract

SARS-CoV-2 has infected over 160 million people and resulted in more than 3.3 million deaths, and we still face many challenges in the rollout of vaccines. Here, we use the high-density microarray patch to deliver a SARS-CoV-2 spike subunit vaccine directly to the skin. We show the vaccine, dry-coated on the patch is thermostable, and delivery of spike via HD-MAP induced greater cellular and antibody immune responses, with serum able to potently neutralize clinically relevant isolates including those from the B.1.1.7 and B.1.351 lineages. Finally, a single dose of HD-MAP-delivered spike provided complete protection from a lethal virus challenge, demonstrating that HD-MAP delivery of a SARS-CoV-2 vaccine is superior to traditional needle-and-syringe vaccination and has the potential to greatly impact the ongoing COVID-19 pandemic.

## Main Text

SARS-CoV-2 is a novel betacoronavirus that emerged in 2019 in Wuhan, China (*1*), before rapidly spreading around the globe causing the COVID-19 pandemic. As of the 19^th^ of May 2021, SARS-CoV-2 has caused over 163 million infections resulting in over 3.3 million deaths (https://covid19.who.int). The rapid spread of SARS-CoV-2 was accompanied by the rapid development of multiple vaccines. Many vaccine modalities were trialed including mRNA (*2–4*), DNA (*5*), protein subunit (*6, 7*), viral vectored (*8–11*), nanoparticle (*12*) and inactivated virus (*13, 14*), and several have subsequently been granted emergency authorization for use in humans. However, the emergence of SARS-CoV-2 variants of concern which possess varying degrees of resistance to neutralization by sera from vaccinated individuals is raising concerns of vaccine escape, highlighting the need for further vaccine research and development (*15–18*). Furthermore, some of these vaccines require ultra-low temperature storage and show only limited stability at 2-8 °C (*19*), making transport and global distribution challenging, especially to resource limited countries. Meeting the demand of vaccine doses for global vaccination coverage in a pandemic has also proven problematic, and dose-sparing methods could prove useful.

One such approach to improving vaccine stability, dose-sparing and ease of distribution is the high-density microarray patch (HD-MAP). The HD-MAP is a 1 x 1 cm solid polymer microprojection array containing 5,000 projections of 250 μm in length (*20*). Vaccine is coated onto the microprojections via a nitrogen-jet-based drying process (*20*) before application to the skin at a velocity of 18-20 m/s via a spring-loaded applicator. This delivers the vaccine directly to the dermal and upper dermal layers of the skin, which are rich in antigen-presenting cells (*21–23*). This vaccine modality offers considerable advantages in thermostability and antigenicity (*20, 24*) with dose-sparing of a large range of vaccines previously observed (*25–32*). Additionally, HD-MAP application avoids the generation of sharps waste and offers advantages in terms of ease of application, potentially negating the need for highly trained healthcare workers (*33, 34*).

Here, we explore HD-MAP delivery of a subunit vaccine candidate – a recombinant SARS-CoV-2 spike glycoprotein, termed HexaPro which has been stabilized in its prefusion conformation by removal of the furin cleavage site and the inclusion of six stabilizing proline mutations(*35*). We found that immunization of mice with spike via HD-MAP application induced significantly higher antibody levels than intradermal needle-and-syringe delivery, with neutralization of virus seen after just one dose. Additionally, immunization of transgenic mice expressing human angiotensin-converting enzyme 2 (ACE2) with spike protein via HD-MAP delivery and adjuvanted with QS21 provided complete protection from a lethal SARS-CoV-2 challenge after just one dose, with no virus replication observed in the lungs or brain of the mice.

## Results

### SARS-CoV-2 Spike (HexaPro) expression and formulation development

HexaPro (*35*), a stabilized SARS-CoV-2 spike protein containing six proline substitutions and a mutated furin cleavage site (hereafter referred to as spike) was recombinantly expressed using the Expi293-F™ expression system and purified via immunoaffinity chromatography. After initial *in vitro* characterization to confirm integrity and conformational folding of the protein (Fig. S1), we investigated formulation and coating optimization on the HD-MAP (Fig. 1A). Initial experiments focused on the selection of generally regarded as safe (GRAS) excipients that stabilize the protein in both the dry-down process and storage over time. Excipients tested included sugars, proteins and denaturing agents, which have previously been shown to stabilize vaccines on the HD-MAP (*27, 29*). Stability was assessed by ELISA, through the binding of a conformational dependent, spike-reactive antibody S309 (*36*). While minimal reduction in antigenicity was seen during the dry down process (measured following immediate reconstitution), storage for 1 or 7 days at 4 °C resulted in considerable loss of antigenic reactivity for some excipients (Fig. S2). However, all formulations containing human serum albumin (HSA) resulted in antigenic stability at all timepoints.

**Fig. 1.**
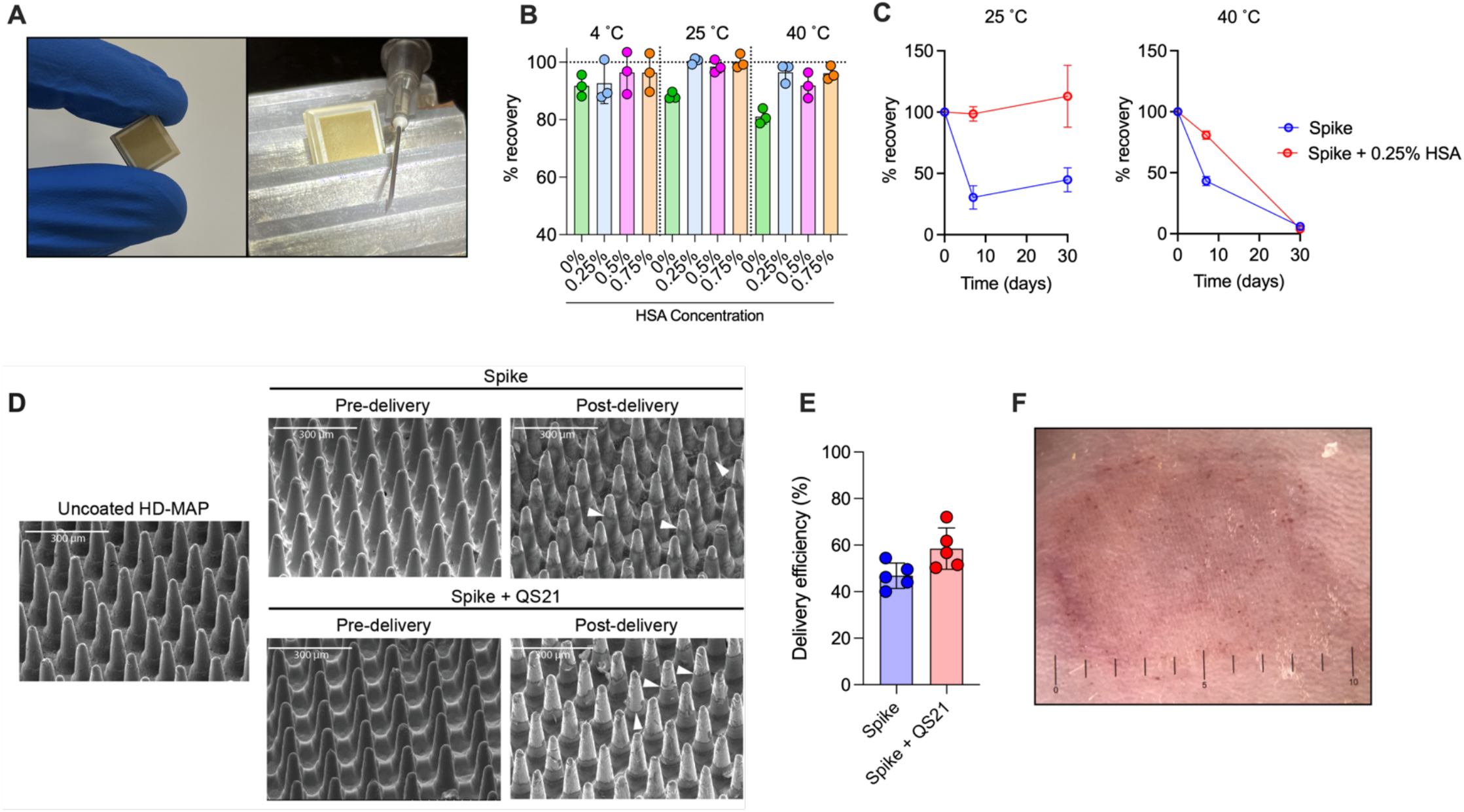
Spike vaccine and the HD-MAP. **(A)** The HD-MAP relative and 27G needle. Temperature stability of dry-coated spike protein with various concentrations of human serum albumin (HSA) after seven days at the indicated temperatures **(B)** or with 0.25% HSA for the indicated temperatures and time **(C).** Data presented as mean with error bars representing SEM. **(D)** scanning electron microscope images of HD-MAPs either uncoated or coated with spike vaccine formulations, pre- and post-delivery into the skin of mice. White arrowheads indicate levels of vaccine removal post-delivery. **(E)** Delivery efficiency into the skin of mice of spike and spike + QS21-coated HD-MAPs. **(F)** HD-MAP delivery site on the flank of a mouse immediately after HD-MAP application and removal.

Next, we investigated the stability of spike-HSA formulations at elevated temperatures and to further optimize the concentration of HSA required to confer stability. To achieve this, spike was dried with or without HSA (0.25 – 0.75% w/v) and stored at 25 or 40 °C and assessed for antigenic stability by ELISA as before. The losses upon storage at both 25 and 40 °C ranged between 1 and 13% after 1 week when HSA was included in the formulation, with higher losses in antigenic integrity of more than 20% at 40 °C observed with spike alone (Fig. 1B). Given the minimal differences in stabilization afforded by all HSA concentrations tested, we chose to proceed with the lowest effective concentration of 0.25% for future formulations to minimize the solid mass required to be dried down on the patch projections. The effects of storage over an extended time period was also assessed. After 1 month at 25 °C in the presence of 0.25% HSA, full antigenic reactivity was recovered, suggesting excellent stability (Fig. 1C). While one week of storage at 40 °C resulted in minimal antigenic loss, 1 month storage at this elevated temperature resulted in limited antigenic recovery (Fig. 1C). As the addition of HSA greatly improved antigenic stability at all temperatures tested, HSA at 0.25% was included in all subsequent HD-MAP vaccine formulations.

We next assessed the delivery efficiency of the vaccine from the coated HD-MAP to mouse skin. When applying spike-coated HD-MAPs to the skin of mice, removal of the vaccine coating can be seen by scanning electron microscopy (SEM) of the patches pre- and post-delivery (Fig. 1D). We also included the saponin-based adjuvant QS21, based on previous data demonstrating its potent adjuvanting effects and compatibility for coating onto the HD-MAP(*29, 37*). Efficient removal of vaccines containing QS21 into the skin of mice was also observed. This vaccine removal corresponded to an average delivery of 50-60% of the payload as assessed via ELISA of spike able to be eluted off the patch post-application (Fig. 1E) and was accompanied by visual signs (erythema and petechiae) of HD-MAP engagement with the depilated skin of the animal (Fig. 1F).

### Immunogenicity of HD-MAP delivered spike

We next sought to assess the immunogenicity of SARS-CoV-2 spike delivered with or without adjuvant, and either via HD-MAP application or intradermal (i.d.) injection using needle-and-syringe (N&S). We chose intradermal, rather than intramuscular vaccination as a more relevant comparator to HD-MAP delivery of vaccines to the immune cell-rich environment of the skin. BALB/c mice were immunized twice at 21-day intervals with 2 μg of spike alone or with the addition of QS21 (Fig. 2A). HD-MAPs coated with excipients only were included as controls. Blood was collected 20 days after each immunization and the serum assessed for spike-specific IgG by ELISA. After one immunization, both unadjuvanted and QS21-adjuvanted spike delivered by HD-MAP induced significantly higher (*p* < 0.0001) IgG levels compared to their i.d.-delivered controls (Fig. 2B). This trend was mirrored in the response after 2 doses, with levels approximately 1.6-fold higher after a second immunization in both HD-MAP groups. Promisingly, serum from the unadjuvanted HD-MAP spike group had significantly higher (*p* < 0.0001) IgG titers compared to QS21-adjuvanted i.d.-delivered spike, highlighting the immune-enhancing attributes of the HD-MAP alone. Serum from day 41 also contained high levels of IgG able to bind to the receptor-binding domain (RBD) and N-terminal domain (NTD) of the spike protein (Fig. S3), indicating broad coverage of epitopes known to be bound by neutralizing antibodies (*36, 38–40*). Again, IgG levels were higher in HD-MAP immunized animals compared to their i.d. counterparts. This was especially apparent for NTD-specific IgG levels, where complete seroconversion in both unadjuvanted and adjuvanted groups was seen in HD-MAP groups compared to only partial seroconversion for i.d.-delivered spike (0% for unadjuvanted and 62.5% for adjuvanted).

**Fig. 2.**
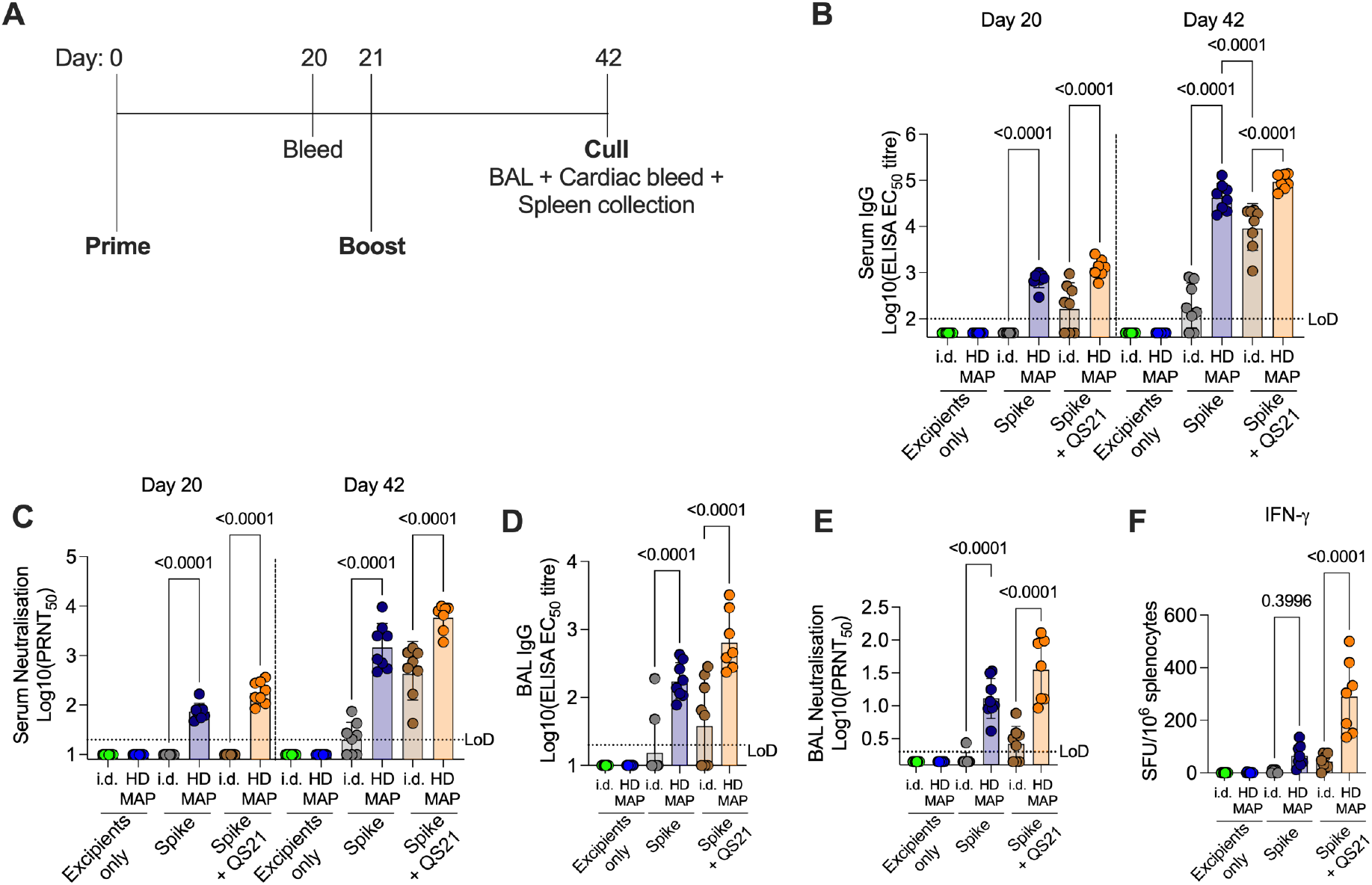
Immunogenicity of HD-MAP spike vaccination in mice. **(A)** Vaccination and bleed schedule of BALB/c mice (n=7 or 8/group) immunized with spike vaccine via HD-MAP application or intradermal delivery. Serum was collected on day 20 or 42 and analysed for **(B)** IgG titer by ELISA and **(C)** virus neutralisation by plaque reduction neutralisation test (PRNT) against the SARS-CoV-2 virus hCoV-19/Australia/QLD02/2020 (GISAID Accession ID EPI_ISL_407896). Bronchoalveolar lavage (BAL) was collected on day 42 and analysed for **(D)** IgG titers as measured by ELISA and **(E)** virus neutralisation by PRNT. **(F)** Interferon-γ ELISpot of splenocytes collected on day 42. Data representative of geometric mean with error bars representing SD. *P* values indicate results of one-way ANOVA with Tukey’s multiple comparison *post-hoc* test.

To investigate the functionality of the induced immune response, we assessed the serum for its ability to neutralize SARS-CoV-2 virus via plaque-reduction neutralization test (PRNT) against an early SARS-CoV-2 isolate, QLD02, matching closely to the original Wuhan-Hu-1 strain. Following one dose, HD-MAP-spike immunized mice had detectable levels of neutralizing antibodies (nAbs) in the serum, which were boosted ~1.7-fold after a second immunization (Fig. 2C). Conversely, spike i.d.-immunize mice had no detectable nAbs after one dose. Although the second i.d. immunization induced a nAb response in mice receiving spike adjuvanted with QS21, this response was significantly lower than their HD-MAP counterparts (Fig. 2C, *p* < 0.0001).Furthermore, with no adjuvant and after two doses, i.d.-delivered spike induced a nAb response in only a subset (62.5%) of animals.

An important site of immunity for respiratory infections such as SARS-CoV-2 is at mucosal surfaces (*41*) of the respiratory tract, and an ideal vaccine would induce immunity at these surfaces. To assess antibody levels at mucosal surfaces, we performed bronchoalveolar lavage (BAL) at the lungs of mice and analyzed antibody levels via ELISA and PRNT. Similar to trends observed in serum IgG analysis, mice receiving spike via HD-MAP induced significantly higher IgG titers compared to their i.d. counterparts (Fig. 2D, *p* < 0.0001 for both adjuvanted and unadjuvanted groups). Furthermore, all HD-MAP-immunized mice produced virus neutralizing responses, with the inclusion of QS21 boosting this response (Fig. 2E). For i.d.-delivered spike alone, BAL neutralization activity was detected in only one animal. The inclusion of QS21 for i.d. delivery resulted in improved BAL neutralization activity, with 62.5% of mice producing a neutralizing response (Fig. 2E).

Cellular immunity to SARS-CoV-2 is an important component of patient recovery from COVID-19 and in providing protection from viral variants (*42–44*). To provide an indicative measure of the cellular response induced by immunization with HD-MAP delivered spike, spleens from mice were harvested at day 42 and splenocytes assessed via interferon-g (IFNg) ELISpot. Delivery of spike + QS21 by HD-MAP induced an average of 290 spot-forming units (SFU) per 10^6^ splenocytes, which was significantly higher compared to i.d. delivery of the same formulation (*p* < 0.0001, Fig. 2F). With no adjuvant, an average of 65 SFU per 10^6^ splenocytes was seen when delivered via HD-MAP, compared to only 6 for i.d. delivery. This is a good initial indication that cellular immunity is induced by HD-MAP immunization, which complements the high IgG titers in the serum and at mucosal sites.

Serum was further analyzed for neutralization of SARS-CoV-2 variants of concern that have recently emerged (*15–18*). These variants, which included an isolate containing the D614G mutation in the spike protein (*45*), a lineage B.1.1.7 isolate and a lineage B.1.351 isolate, have been shown to have a functional phenotype characterized by increased ACE2 binding affinity, increased transmissibility, and escape from neutralizing antibody binding (*15–18, 45, 46*). Promisingly, nAb levels in day 42 serum from HD-MAP-immunized mice showed no significant decrease in neutralizing activity when tested against any of these variants, although there was a trend for decreased responses against B.1.351 (Fig. 3, Extended data/Supp. Fig. S4). When spike was delivered via the i.d. route, significant decreases in nAb activity (*p* = 0.0159) were seen against the B.1.351 lineage virus, though all samples were still neutralizing (Fig. 3B).

**Fig. 3.**
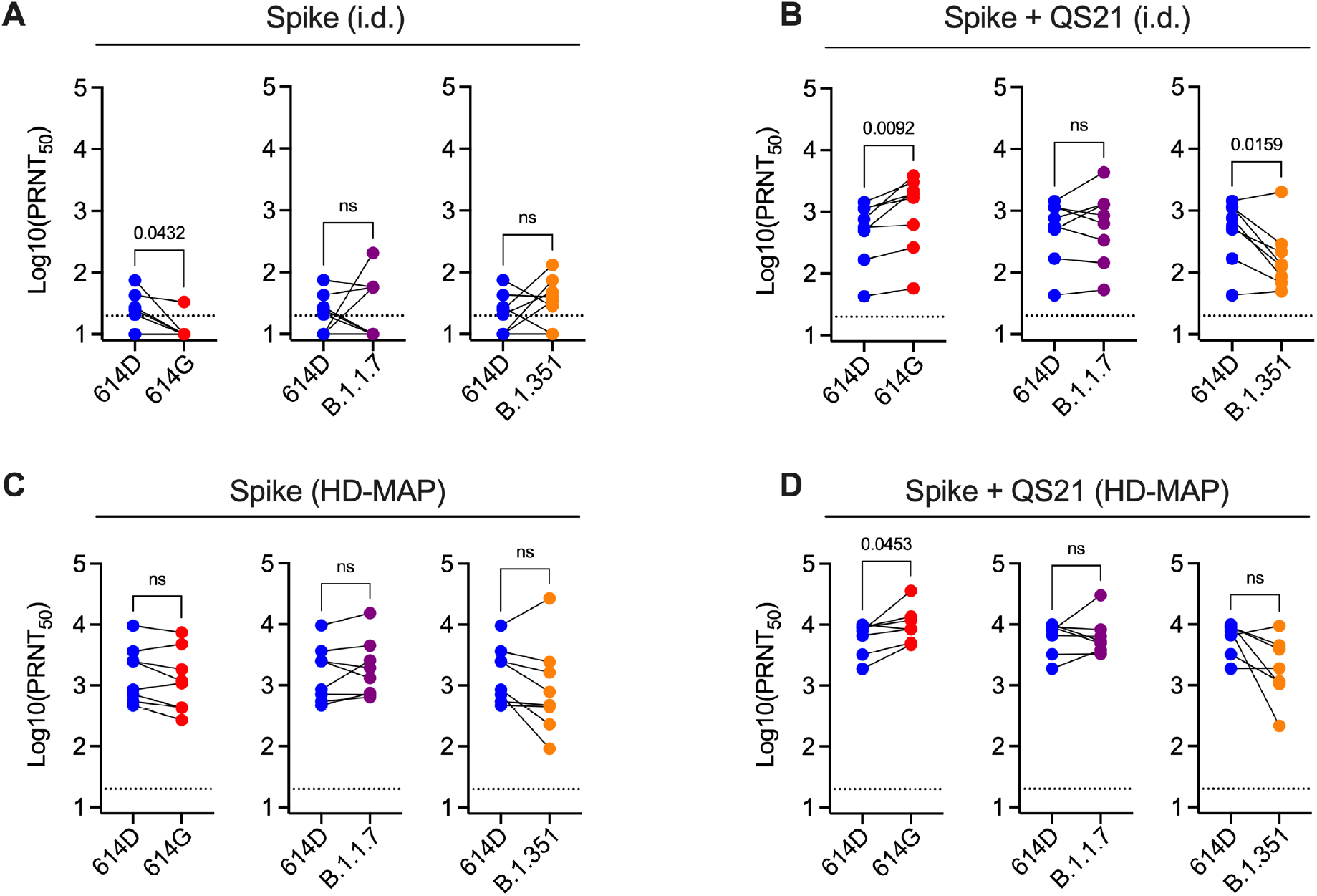
Serum neutralization against SARS-CoV-2 viral variants. Serum from mice (n=7 or 8/group) was collected 21 days after the second immunization with SARS-CoV-2 spike with or without the adjuvant QS21, either via i.d. injection (A, B) or HD-MAP application (C, D). Serum was tested for neutralization against an isolate containing the 614G mutation (hCoV-19/Australia/QLDID935/2020, GISAID Accession ID EPI_ISL_436097), a lineage B.1.1.7 virus (hCoV-19/Australia/QLD1517/2021, GISAID Accession ID EPI_ISL_944644), and a lineage B.1.351 virus (hCoV-19/Australia/QLD1520/2020, GISAID Accession ID EPI_ISL_968081), and PRNT50 values compared to the parental strain containing the 614D in the spike protein (hCoV-19/Australia/QLD02/2020, GISAID Accession ID EPI_ISL_407896). Data represents geometric mean of individual mice and *P* values represent results of a paired two-tailed t-test. Dotted lines show assay limit of detection.

### Protection in mice after a single dose of HD-MAP delivered spike

With the observation that HD-MAP immunization induced significantly higher IgG and cellular immune responses relative to N&S i.d. delivery, we next assessed HD-MAP vaccine potency in a SARS-CoV-2 challenge model. Transgenic mice expressing human ACE2 driven by the cytokeratin-18 (K18) promoter (K18-hACE2 mice), have been shown to be an effective model for SARS-CoV-2 infection (*47*) and were used in this study. Mice (n=8) were immunized via HD-MAP with either a single dose or a prime/boost regime at 21-day intervals, with or without QS21 as an adjuvant, or left unimmunized as naïve controls (Fig. 4A). Blood was taken for analysis of serum IgG levels via ELISA and PRNT prior to challenge, which occurred 21 days after the final immunization. At this time, all immunized mice had seroconverted with high levels of spike-specific IgG, and all had high levels of nAbs against 614D, 614G and B.1.1.7 virus isolates, though nAb levels against the B.1.351 isolate were lower (Fig. 4B, Fig S5). Mice were then challenged with 10^4^ PFU of an early isolate of SARS-CoV-2 (hCoV-19/Australia/VIC01/2020, GISAID Accession ID EPI_ISL_406844, matching to the Wuhan-1 reference virus (*48*)) via intranasal inoculation, and monitored daily for weight loss and clinical signs of infection. By day 5 postinfection, all naïve mice were showing clinical signs of infection, and subsequently required euthanasia by day 6 due to weight loss and clinical scores (Fig. 4C, D and E). Of those mice receiving a single dose of unadjuvanted spike, 4 out of 8 also showed clinical signs of infection and required euthanasia at day 6. All other immunized mice had no observable signs of clinical infection or weight loss at this time. To compare viral loads and histopathology of tissue at the peak of infection, 4 out of 8 animals from all other groups were also sacrificed. The remaining mice were then monitored daily until day 14 post-infection. Both the lungs and brains of naïve mice showed high levels of virus at day 6 post-infection (log-transformed mean titers of 4.2 PFU/g in the lungs and 8.8 PFU/g in the brains) (Fig. 4F and G), consistent with previous reports of SARS-CoV-2 infection in K18-hACE2 mice(*47, 49*). The mice receiving a single dose of unadjuvanted spike had detectable virus levels in the brain (log-transformed titer of 6.8 PFU/g), albeit at significantly lower levels (*p* = 0.0005) than naïve mice, but no detectable virus in the lungs (Fig. 4F and G). These mice had however been sacrificed due to signs of infection. Mice in this group that did not show symptoms were not analyzed at this timepoint. Conversely, all other immunization regimes (a single dose of QS21-adjuvanted spike, two doses of adjuvanted spike and two doses of unadjuvanted spike) completely inhibited infection as evidenced by no detectable virus in the lungs at day 6 post-infection (Fig. 4F and G). Histopathological analysis of lung and brain tissue revealed mild to moderate perivascular and interstitial leukocyte infiltration in the lungs of naïve and single-dose immunized mice (Fig. S6A). This was reduced in mice receiving two doses of vaccine. Similar trends were seen in brain tissue, with perivascular cuffing, neutrophil infiltration, neural degeneration and gliosis seen in naïve and single-dose mice, with mice from two dose groups appearing normal (Fig. S6B).

**Fig. 4.**
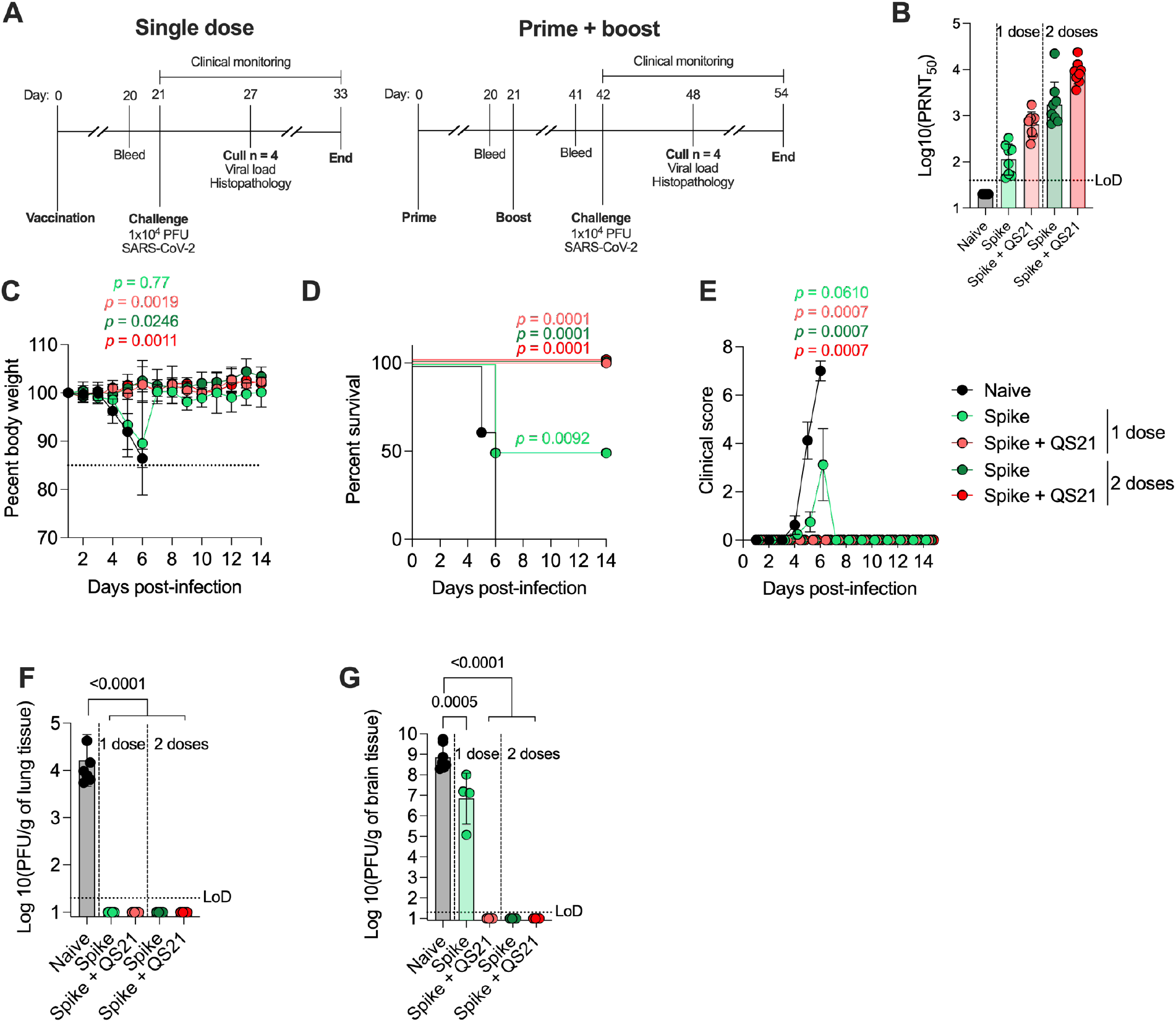
HD-MAP vaccination of K18-hACE2 mice and protection from virus challenge. **(A)** Immunization and challenge schedule for K18-hACE2 mice (n=8/group) vaccinated with HD-MAP spike (single dose or prime/boost regime) and challenge with SARS-CoV-2 (hCoV-19/Australia/VIC01/2020, GISAID Accession ID EPI_ISL_406844). **(B)** Neutralization titres of serum collected prior to challenge (day 20 or 41 for single dose and prime/boost, respectively) against hCoV-19/Australia/QLD02/2020. **(C-E)** Body weight, survival and clinical score of mice throughout the course of the challenge. Viral titer in the **(F)** lungs and **(G)** brain of mice at day 6 post-infection. *P*-values for survival curve represent results of the log-rank test relative to naïve mice with a K value of 4 and a Bonferroni threshold of 0.0125. *P*-values for body weight and clinical score graphs represent results of repeated measures one-way ANOVA relative to naïve mice with Dunnett’s test for multiple comparisons. All other graphs have data representing geometric mean with error bars representing geometric SD and *P*-values indicating results of one-way ANOVA with Tukey’s multiple comparison *post-hoc* test.

## Discussion

Here we report the combination of a SARS-CoV-2 spike subunit protein vaccine candidate with the HD-MAP vaccine delivery platform, resulting in a stable and effective vaccine candidate against SARS-CoV-2. With the HD-MAP platform, dry-coated vaccine is delivered directly to the immune-rich epidermis and upper dermal layers of the skin, while simultaneously causing localized cell death and inflammation, both of which serve to enhance vaccine-induced immunity (*22, 50*). This platform delivery system offers a number of other benefits, including ease of administration, and potentially self-administration or administration by individuals who are not trained health-care professionals, reduced cold chain dependence, no requirement for reconstitution and improved thermostability. We showed that the dry-coated spike protein was thermostable (as measured by conformational epitope integrity), with stability observed for up to a month at 25 °C and a week at 40 °C (Fig 1B and C). Should this measure of stability translate to biophysical stability, this represents a marked improvement over other SARS-CoV-2 vaccine candidates, where limited stability (2-24 hours) at room temperature is seen (*19*). This lends itself well to improvements in vaccine transport and delivery to patients, especially in a “last-mile” context where appropriate infrastructure may be limited. This is particularly relevant to low-to-middle income countries (LMICs), where there is an urgent need to facilitate vaccination of populations against SARS-CoV-2 (*51, 52*).

HD-MAP-delivered spike resulted in a robust neutralizing IgG and cellular immune response in mice, which was superior to delivery of the same vaccine via intradermal needle-and-syringe immunization. This is consistent with previous studies utilizing the HD-MAP where significant improvements in immunogenicity were observed (*27, 29, 53, 54*). The inclusion of QS21, which is known to broaden the immune response and enhance cell-mediated immunity (*37*), induced high levels of spike-specific **γ**-IFN-secreting T cells and increased the magnitude of RBD- and NTD-specific antibodies. The broad IgG response observed, resulted in neutralization of representative emerging viral variants, including an isolate containing the 614G mutation in the spike protein and isolates from both the B.1.1.7 and B.1.351 lineages. These variants, which appear to have supplanted the ancestral virus, have shown escape from neutralization by some monoclonal nAbs and vaccine-induced serum (*15–18, 45, 46*). These collective findings are promising given the importance of cellular immunity in SARS-CoV-2 disease resolution and protection from emerging viral variants (*42, 44*). Additionally, the HD-MAP appears to provide improved immunogenicity and cellular responses to SARS-CoV-2 vaccine candidates when compared to immunization with other lower density microarray patch delivery systems (*55, 56*). Taken together, this data suggests HD-MAP vaccination could provide improved cellular immune responses and increased breadth of IgG, both of which would limit the potential for escape from protection by emerging viral variants.

Immunization of K18-hACE2 mice with spike via HD-MAP resulted in complete protection from disease after just a single QS21-adjuvanted dose. The protective immune response resulted in abrogation of virus replication in both the lungs and brain tissue assessed on day 6 post-infection, in contrast to naïve mice which had high levels of replicating virus in these organs (Fig. 4F and G). This is notable as SARS-CoV-2 infection has been shown to infect the brain of humans and mice, causing meningitis and encephalitis (*49, 57, 58*). Protection with a single dose would provide significant benefits for rollout of mass vaccination campaigns during a pandemic. Only requiring a single dose to achieve protection would mean fewer doses to be manufactured and more rapid vaccine coverage of populations, in contrast to most current SARS-CoV-2 vaccines which require 2 doses 3-12 weeks apart to achieve protection (*59–62*). Although some adenovirus-vectored vaccines have been shown to provide single-dose protection from SARS-CoV-2 infection in mice and humans (*63, 64*), issues surrounding safety and undesired adverse events have impacted their clinical rollout (*65*). Subunit vaccines could offer an improved and more desirable safety profile.

To date, only limited studies investigating subunit-based SARS-CoV-2 vaccines have been performed, and while single-dose protection has been seen, this was achieved with much higher doses than the 2 μg used here. Tian *et al* used 10 μg of a full-length spike protein vaccine with Matrix-M adjuvant, and saw protection after a single dose, though signs of infection were still observed in mice (*6*). Another study utilized a spike nanoparticle vaccine in which multiple spike proteins are assembled on a nanoparticle (*66*). Immunization provided protection in hamsters, though this immunity was not complete and required a dose of 100 μg, 50 times higher than required by HD-MAP immunization (*66*). Other work using a prefusion-stabilized spike protein (5 μg/dose, adjuvanted with MF59) showed reduced viral load in the lungs and throat swabs of hamsters on day 4 post-infection after one or two immunizations (*7*), with relatively high levels of replicating virus still recovered from these samples. Additionally, neither regime decreased the viral load in the nasal turbinates compared to naïve hamsters on day 4 or day 8 post-infection (*7*). This suggests the antibody and/or cellular immunity induced by these vaccination regimes were not high enough to inhibit viral replication in these organs.

In conclusion, this body of work demonstrates the first investigation of SARS-CoV-2 spike vaccination via a microarray patch. HD-MAP spike vaccines are stable, immunogenic, and protective against virus challenge in mice after a single dose. These finding represent a substantial improvement in many areas of SARS-CoV-2 vaccination and offers a promising alternative to currently available vaccines that warrants further investigation in the context of human SARS-CoV-2 infection.

## Materials and Methods

### Animal ethics and animals

Female BALB/c and K18-hACE2 transgenic mice were purchased from the Animal Resources Centre (Perth, Australia) and housed in pathogen-free conditions at the Australian Institute for Bioengineering and Nanotechnology or Griffith University. All animal procedures were approved by the University of Queensland and Griffith University animal ethics committees (approval numbers: SCMB/322/19/AIBN and MHIQ/12/20/AEC).

### Cell lines

Expi293-F™ and ExpiCHO-S™ cells were purchased from ThermoFisher Scientific and maintained as per the manufacturer’s instructions. VeroE6 cells were purchased from the American Type Culture Collection (ATCC) (catalog: ATCC CRL-1586) and cultured in Dulbecco’s Modified Eagle Medium (DMEM) supplicated with 10% fetal bovine serum (Bovogen, USA), penicillin (100 U/mL) and streptomycin (100 μg/mL) at 37 °C with 5% CO_2_.

### Viruses

Low passage virus isolates of SARS-CoV-2 that were recovered from nasopharyngeal aspirates of infected individuals were kindly provided by the Queensland Health Forensic and Scientific Services, Queensland Department of Health, Peter Doherty Institute for Infection and Immunity and Melbourne Health, Victoria, Australia. Virus isolates used in this study include the early Australian isolates hCoV-19/Australia/QLD02/2020 (GISAID Accession ID EPI_ISL_407896, collected on the 30^th^ of January 2020) and hCoV-19/Australia/VIC01/2020(*48*) (GISAID Accession ID EPI_ISL_406844, collected on the 25^th^ of January 2020), an isolate containing the D614G mutation (hCoV-19/Australia/QLDID935/2020, GISAID Accession ID EPI_ISL_436097, collected on 25^th^ March 2020), an isolate of the B.1.1.7 lineage (hCoV-19/Australia/QLD1517/2021, GISAID Accession ID EPI_ISL_944644, collected on 6^th^ of January 2021), and an isolate of the B.1.351 lineage (hCoV-19/Australia/QLD1520/2020, GISAID Accession ID EPI_ISL_968081, collected on 29^th^ December 2020). Virus isolates were further propagated on VeroE6 cells and stocks stored at −80 °C. Virus titer was determined by immunoplaque assay as previously described (*67*).

### Plasmids

The plasmid encoding SARS-CoV-2 S HexaPro was a gift from Jason McLellan (Addgene plasmid # 154754; http://n2t.net/addgene:154754; RRID:Addgene_154754). This construct has been modified to include a cleavage site substitution as well as six prolines for stability (*35*). Plasmids encoding heavy and light chains of SARS-CoV-2 S-specific antibodies, as well as spike receptor binding domain and N-terminal domain constructs were constructed in-house as previously described (*7, 67, 68*).

### Recombinant antibody expression and purification

Plasmids encoding the heavy and light chains of monoclonal antibodies were prepared using the PureYield™ Plasmid Midi- or Maxiprep system and filter sterilized using a 0.22 μm filter. Plasmids were then transfected into ExpiCHO-S™ cells (ThermoFisher Scientific) as per the manufacturer’s instructions. Culture supernatant was harvest via centrifugation at 7 days post-transfection, and antibodies purified using Protein A affinity chromatography on an AKTA pure FPLC system (Cytiva) as per the manufacturer’s instructions. Purified antibodies were concentrated and buffer exchanged into phosphate-buffered saline (PBS) using a 30 kDa MWCO Amicon Ultra Centrifugal Filter (Merk) and stored at −20 °C until use.

### Fc-fusion protein expression and purification

Plasmids containing the receptor binding domain (RBD) and N-terminal domain (NTD) of SARS-CoV-2 spike fused to human Fc were prepared using the PureYield™ Plasmid Midi- or Maxi-prep system and filter sterilized using a 0.22 μm filter. Plasmids were then transfected into ExpiCHO-S™ cells (ThermoFisher Scientific) as per the manufacturer’s instructions. Culture supernatant was harvested and protein purified via Protein A affinity chromatography as before.

### SARS-CoV-2 Spike HexaPro expression and purification

Plasmid DNA for transfection was prepared using Promega PureYield™ Plasmid Maxiprep System and filter-sterilised using a 0.22 μm filter prior to transfection into Expi293-F™ cells (ThermoFisher Scientific), as per the manufacturer’s instructions. Culture supernatant was clarified by centrifugation at 5,000 x g for 30 minutes at 4 °C before filtration through a 0.22 μm filter. Protein was purified from supernatants as previously described(*7*), using an in-house made immunoaffinity purification column containing the S-specific mAb 2M-10B11. After purification, proteins were concentrated and buffer exchanged into PBS as before, and stored at −20 °C. Protein was assessed for purity SDS-PAGE using a 4-12% NuPage SDS gel and size-exclusion chromatography using a Superdex 200 Increase 10/300 GL column. Purified spike protein was also analyzed via ELISA. Briefly, SARS-CoV-2 S protein was coated onto ELISA plates at 2 μg/mL in PBS and incubated at 4 °C overnight. The next day, plates were blocked with 5% milk diluent blocking concentrate (Seracare) for 30 minutes at room temperature. Serial dilutions of antibodies were then added to the plates before incubation at 37 °C for 1 hour. After washing with PBS with 0.05% tween 20, relevant HRP-conjugated secondary antibodies were added (goat anti-mouse or anti-human, diluted 1:2000). After washing, TMB substrate was then added to develop the ELISA prior to stopping the reaction with 1M phosphoric acid. Absorbance was then read at 450 nm using a Varioskan LUX Microplate reader (ThermoFisher).

### Stability dry down assay

Spike protein was formulated as desired and added to wells of a 96-well tissue culture plate before being placed under a filtered nitrogen gas jet stream at 15 liters per minute for 15 minutes using a MICROVAP microplate evaporator (PM Separations). Plates were then sealed in foil bags containing desiccant and stored for the desired time. To reconstitute, 150 μL of 5% Milk diluent blocking concentrate (Seracare) diluted in PBS containing 0.05% tween 20 was added to each well. Reconstitution was achieved by pipetting and incubating for 15 minutes at 37 °C with shaking at 125 rpm. Proteins were then analyzed via capture ELISA as above, with samples captured on streptavidin-coated plates and probed with S-specific antibodies.

### Negative stain transmission electron microscopy

Purified spike protein was diluted to ~10 μg/mL in PBS and applied to glow-discharged carbon films supported by formvar on 400-mesh copper grids (ProSciTech). Samples were applied for 2 minutes before washing three times with water. The grids were then stained with 1% uranyl acetate and air dried before imaging on a Hitachi HT7700 transmission electron microscope at 120 kV and 30k magnification.

### High density microarray patch coating and application

High density microarray patches (HD-MAPs) were kindly provided by Vaxxas Pty Ltd (Brisbane, Australia). Prior to vaccine coating, HD-MAPs were cleaned via oxygen plasma treatment for 5 minutes at 30W power at the Australian national Fabrication Facility, Queensland Node. A vaccine coating solution consisting of various amounts of SARS-CoV-2 HexaPro spike protein, 0.25% human serum albumin (HSA), 0.75% methylcellulose, and varying amounts of the adjuvant QS21 (Desert King) was then applied to the patch. The solution was then dried using a sterile-filtered nitrogen gas stream as previously described(*54*). Vaccine-coated HD-MAPs were then applied to the flank of mice using who had previously had the hair at the vaccination site removed via shaving and depilatory cream. The HD-MAPs were applied at a velocity of 18-20 m/s using a custom applicator.

### Scanning electron microscopy of HD-MAPs

HD-MAPs were coated with 15 nm of platinum and imaged by scanning electron microscopy (SEM) using a Hitachi SU3500 at the Centre for Microscopy and Microanalysis at the University of Queensland. Samples were at a 45 °C angle.

### Immunization of BALB/c mice

Naïve 6-8 week old female BALB/c mice were randomly divided into 6 groups of 8 mice and vaccinated either via HD-MAP application or intradermal injection with 2 μg of SARS-CoV-2 S HexaPro, 2 μg of SARS-CoV-2 S HexaPro with the adjuvant QS21 (Desert King), or vehicle only control. All mice were immunized twice at 21-day intervals. Blood was taken prior to each immunization and 21 days after the final immunization via tail bleed or cardiac puncture. A bronchoalveolar lavage (BAL) was also performed at the time of cardiac puncture. The blood was allowed to clot overnight at 4 °C before the serum harvested by centrifugation at 10,000 x g for 10 minutes at 4 °C. Samples were stored at −20 °C until further analysis. The BAL fluid was centrifuged at 1000 x g for 5 minutes to remove debris and the supernatant harvested and stored at −80 °C until further analysis.

### ELISpot

Spleens from mice were collected and processed into single-cell suspensions in RPMI1640 media supplemented with 10% heat-inactivated fetal calf serum and penicillin/streptomycin (R10 media). Red blood cells were lysed using eBioscience™ RBC lysis buffer (ThermoFisher) and resuspended in R10 media to stop the lysis. Splenocytes were counted and 100,000 cells/well added to the plates of the Mouse IFN-g ELISpot^PLUS^ (HRP) kit (MABTECH). Cells were then stimulated for 18 hours at 37 °C with SARS-CoV-2 S PepTivator peptides (Miltenyi Biotec) or media alone as a negative control. Spots were developed as per the manufacturer’s instructions. Spots were quantified by eye and plotted as spot-forming units (SFU) per million cells.

### K18-hACE2 mouse challenge model

Female K18-hACE2 mice (6-8 weeks old) were immunized via HD-MAP application as before or left unimmunized as naïve controls. Mice received 1 or 2 immunizations, 21 days apart, and were challenged with 1 x 10^4^ PFU of SARS-CoV-2 (VIC01 isolate) 21 days after the final immunization via intranasal inoculation (20 μL total volume) while under isoflurane anesthesia. Blood was taken prior to each immunization and prior to challenge. Mice were monitored daily for weight loss and disease severity and culled when weight loss surpassed 15% of pre-challenge body weight. After culling, lung and brain tissue were harvested for viral titer assessment immunoplaque assay. A lung lobe and brain hemisphere from each animal was fixed in 10% neutral buffered formalin and processed for paraffin embedding and hematoxylin-eosin stained sections were examined by a veterinary pathologist blinded to the treatments.

### Serum ELISAs

Serum from mouse immunizations was assessed for SARS-CoV-2 S-specific IgG levels via ELISA as previously described (*7*). Briefly, SARS-CoV-2 spike protein was coated onto ELISA plates at 2 μg/mL in PBS overnight at 4 °C. Plates were blocked and serial dilutions of serum were added to the plates for 1 hour at 37 °C. Serum binding was detected using an HRP-linked goat anti-mouse secondary antibody and TMB substrate as discussed previously.

### Plaque reduction neutralization tests

Serum and BAL fluid was assessed for SARS-CoV-2 neutralization activity via plaque reduction neutralization test (PRNT) as previously described (*67*). Briefly, heat-inactivated serum or BAL fluid were serially diluted in DMEM supplemented with 2% FBS and penicillin/streptomycin (Gibco) before virus was added. Samples were incubated for 30 minutes at 37 °C before being added to confluent VeroE6 monolayers in 96-well plates. Infection was allowed to proceed for 1 hour at 37 °C before virus inoculum removed and overlay medium added. Cells were fixed with 80% acetone 14-16 hours post-infection and allowed to dry prior to plaque visualization with SARS-CoV-2 spike-specific mAbs CR3022 or S309 and IRDye 800CW-conjugated goat anti-human secondary antibodies. Plates were scanned using an Odyssey CLx imaging system (LI-COR).

### Assessment of viral titer in mouse organs

Viral titer in the lungs of mice was determined via immunoplaque assay as previously described (*67*). Briefly, lung and brain tissues were homogenized on a Bead Ruptor 24 Elite (Omni International, Kennesaw, GA) and centrifuged at 12,000 x g for 7.5 minutes at 4 °C before performing serial dilutions, which were added to VeroE6 cells in a 96-well tissue culture plates. Plates were fixed and plaques visualized via immunostaining.

### Statistical analysis

GraphPad Prism 9 software was used for statistical analysis and generation of graphs and figures. Data are presented as mean with standard deviation or standard error in the mean, as described in the figure legends. Statistical significance was determined via one-way ANOVA with Tukey’s multiple comparison test, with p < 0.05 considered statistically significant.

## Acknowledgments

We would like to thank the Queensland Health Forensic and Scientific Services, Queensland Department of Health, as well as the Peter Doherty Institute for Infection and Immunity and Melbourne Health, Victoria, Australia, for providing SARS-CoV-2 isolates. We thank A/Prof. Keith Chappell, Dr Noushin Jaberolansar and Dr Andrew Young from the University of Queensland for providing technical advice and reagents for antigen purification. We thank the staff at the University of Queensland Biological Resources facility for providing technical assistance. We acknowledge the facilities, and the scientific and technical assistance, of the Australian Microscopy and Microanalysis Research Facility at the Centre for Microscopy and Microanalysis, the University of Queensland. We also thank the staff at the Protein Expression Facility, the University of Queensland. J.J.Y.C and A.I. are the recipients of the University of Queensland Research Training Scholarship. This work was supported by Advance Queensland Industry Research Fellowship sponsored by Vaxxas and Technovalia. This work was performed in part at the Queensland Node of the Australian National Fabrication Facility, a company established under the National Collaborative Research Infrastructure Strategy to provide nano- and micro-fabrication for Australia’s researchers. We thank Mr Hamish McMath for assistance with HD-MAP delivery, BL3 facility and Vivarium at Griffith University. We would also like to thank Dr Adam Taylor from the School of Medical Science, Griffith University, for assistance with animal work in the PC3 facility. This work was supported by MRFF 2001931 to N.A.J.M. HD-MAPs were kindly provided by Vaxxas Pty Ltd.

## Data and materials availability

All data are available in the main text or the supplementary materials

