## Supplemental figures for "Complete protection by a single dose skin patch delivered SARS-CoV-2 spike vaccine"

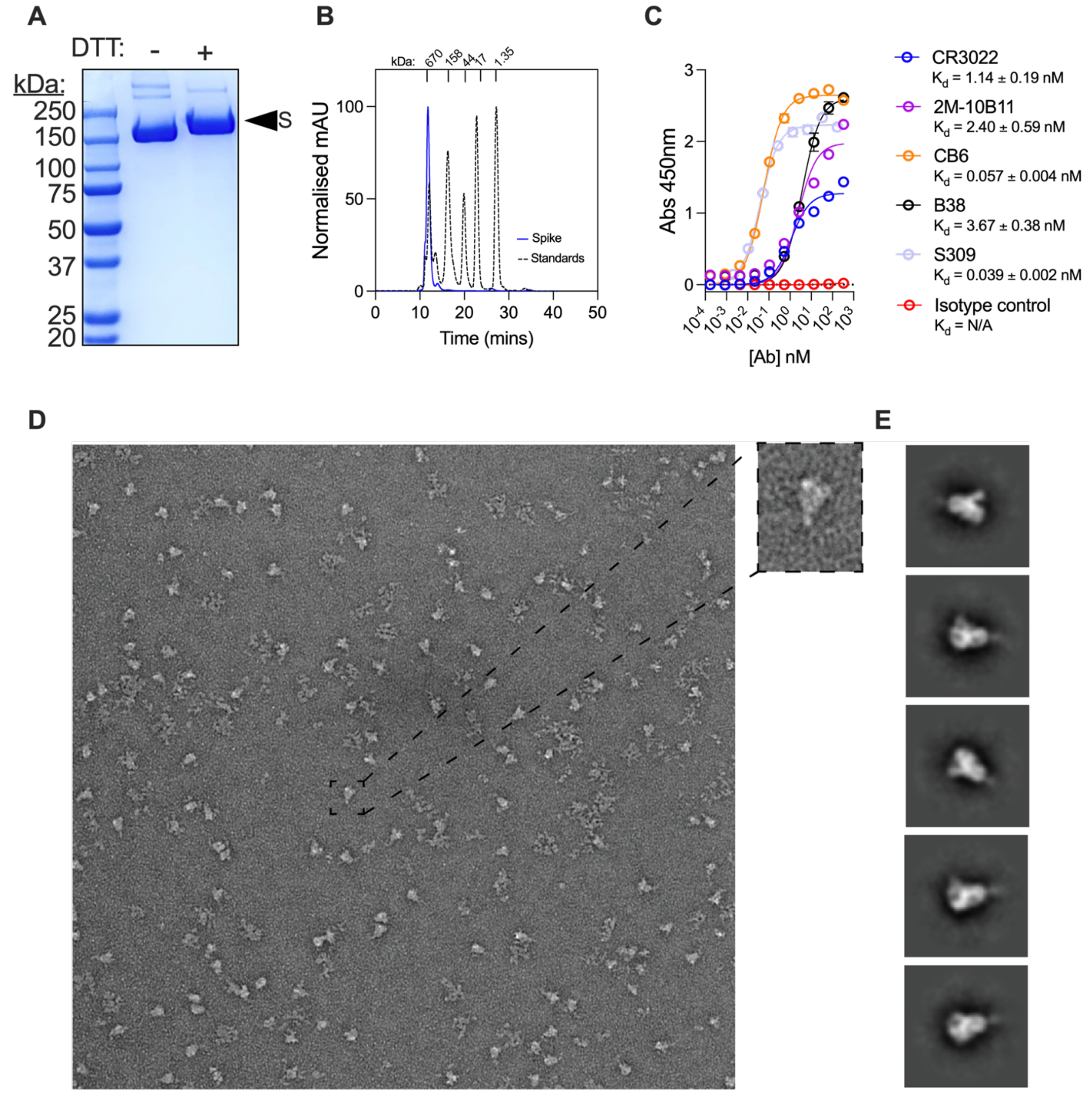


**Fig. S1.** ***In vitro*** **characterisation of SARS-CoV-2 spike. (A)** Coomassie-stained SDS-PAGE of purified SARS-CoV-2 S HexaPro with or without DTT. **(B)** Analytical size-exclusion chromatography of purified HexaPro spike on a Superdex 200 Increase 10/300 GL column, including size markers. **(C)** ELISA of purified HexaPro spike with a panel of SARS-CoV-2 spike-specific monoclonal antibodies. Data represents mean of n=2 technical replicates and error bars represent SD. **(D)** Negative-stain TEM of purified HexaPro spike with representative 2D class averages shown in **(E)** with a box size of 500Å.

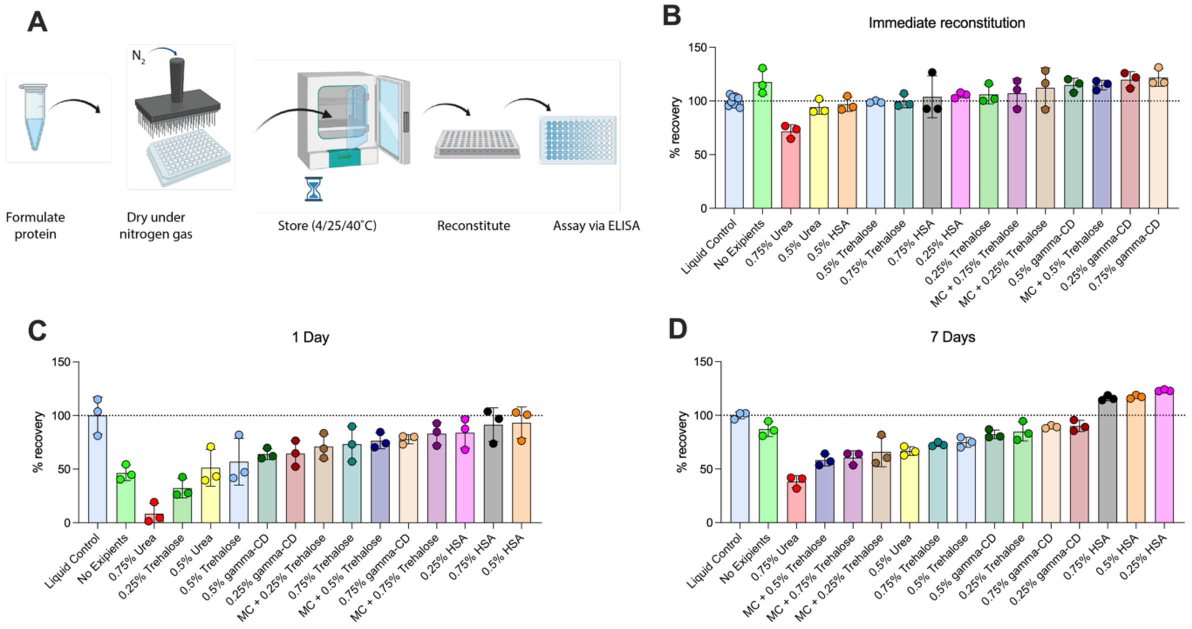


**Fig. S2. Stability of dried spike. (A)** Schematic of the process of excipient screening. SARS-CoV-2 spike was dried in various excipients and reconstituted **(B)** immediately or stored at 4 ˚C for **(C)** 1 or **(D)** 7 days prior to reconstitution. Recovered protein was analysed via ELISA with the conformation dependent S-specific mAb S309. Percent recovery is relative to a liquid control prepared fresh on the day of each assay.

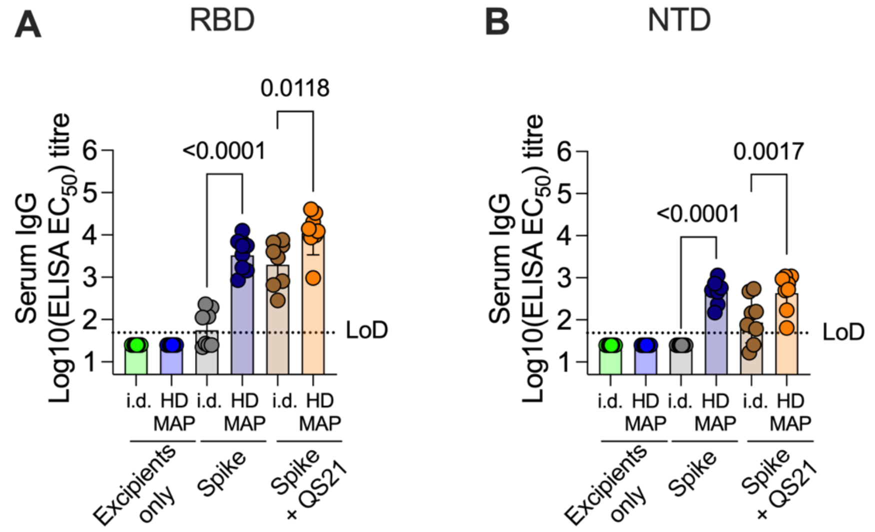


**Fig. S3. Domain-specific IgG titers.** Serum from mice immunized with SARS-CoV-2 spike was assessed by ELISA against **(A)** the receptor binding domain (RBD) or **(B)** N-terminal domain (NTD) of the spike protein. Data representative of geometric mean with error bars representing geometric SD. *P* values indicate results of one-way ANOVA with Tukey's multiple comparison *post-hoc* test.

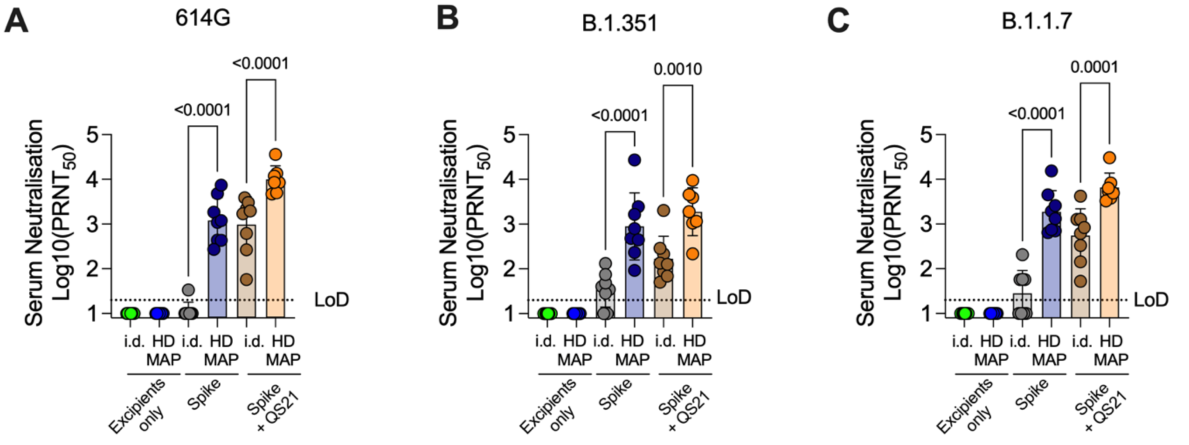


**Fig. S4. Neutralization of SARS-CoV-2 variants.** Serum from mice immunized with SARS-CoV-2 spike via intradermal (i.d.) delivery or HD-MAP delivery, with or without QS21 as an adjuvant, was assessed for neutralization against SARS-CoV-2 variants **(A)** containing the 614G mutation in the spike protein, **(B)** a lineage B.1.351 virus and **(C)** a lineage B1.1.7 virus. Data representative of geometric mean with error bars representing geometric SD. *P* values indicate results of one-way ANOVA with Tukey's multiple comparison *post-hoc* test.

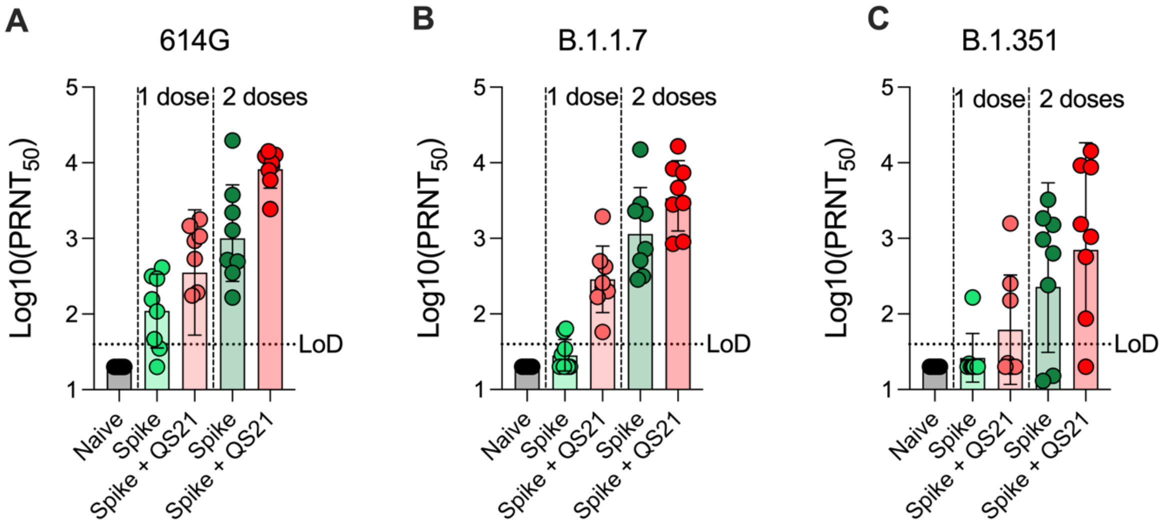


**Fig. S5. SARS-CoV-2 variant neutralization by K18-hACE2 mouse serum.** Serum from K18-hACE2 mice immunized with 1 or 2 doses of HD-MAP delivered spike was analysed for neutralization by PRNT against SARS-CoV-2 variants **(A)** 614G, **(B)** from the B.1.1.7 lineage and (**C)** from the B.1.351 lineage. Data represents geometric mean and error bars represent geometric SD.

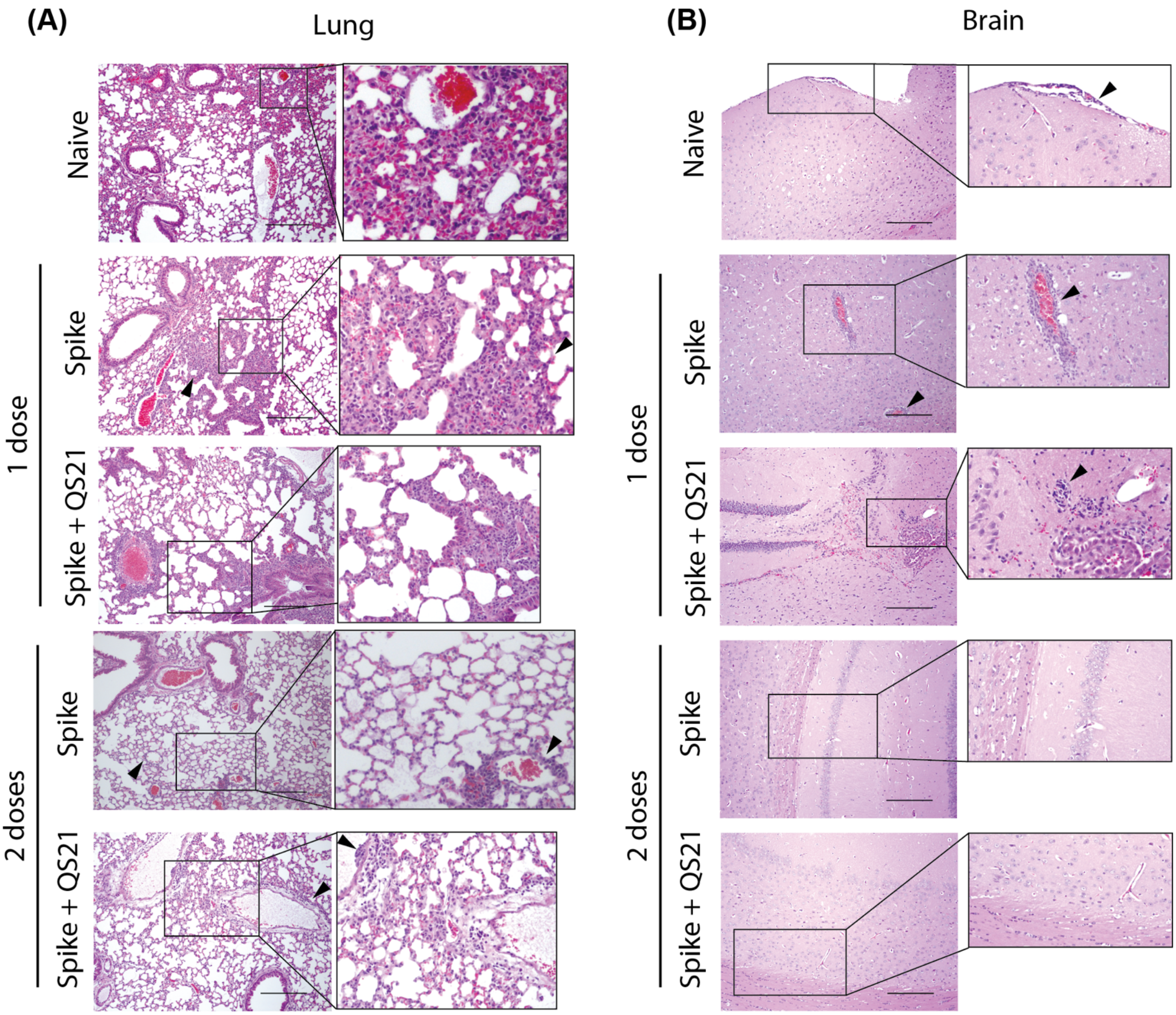


**Fig. S6. Histopathology of lungs and brains from K18-hACE2 challenged with SARS-CoV-2. (A)** Lungs and **(B)** brain tissue from naïve or vaccinated K18-hACE2 mice challenged with SARS-CoV-2 were collected on day 6 post-infection for hematoxylin and eosin staining. Representative images are shown. Arrowheads indicate leukocyte infiltration (lungs) or perivascular cuffing (brain).
